# Cocaine during adolescence differentially impacts psychomotor sensitization and epigenetic profiles in adult male rats with divergent affective phenotypes

**DOI:** 10.1101/2022.07.13.499935

**Authors:** Aram Parsegian, M. Julia García-Fuster, Elaine Hebda-Bauer, Stanley J. Watson, Shelly B. Flagel, Huda Akil

## Abstract

While early life drug use reliably predicts addiction liability in adulthood, the molecular mechanism by which this occurs, and the degree genetic predisposition is involved is not known. We used a unique genetic rat model of temperament to determine the impact of adolescent cocaine experience on adult psychomotor sensitization and epigenetic profiles in the striatum. Relative to adult bred low-responder (bLR) rats, bred high-responders (bHR) are more sensitive to the psychomotor-activating effects of cocaine and reinstate drug-seeking behavior more readily following prolonged cocaine exposure and a period of abstinence. We found that a 7-day sensitizing cocaine regimen (15 mg/kg/day) during either adolescence or adulthood produced psychomotor sensitization in bHRs only, while a dual cocaine exposure prevents further sensitization, suggesting limits on neuroplasticity. By contrast, adolescent cocaine in bLRs shifted their resilient phenotype, rendering them more responsive to cocaine in adulthood when initially challenged with cocaine We then assessed two functionally opposite epigenetic chromatin modifications, previously implicated in addiction liability, permissive acetylation (ac) and repressive tri-methylation (me3), on histone 3 lysine 9 (H3K9) in four sub-regions of the striatum. In bHRs, decreased repressive histone, H3K9me3, and increased permissive histone, acH3K9, in nucleus accumbens core associated with cocaine sensitization. However, there were limits on these epigenetic changes that paralleled the behavioral limits on further sensitization. In bLRs, the combination of cocaine exposure in adolescence and adulthood, which lead to an increase in their responsiveness to a cocaine challenge, also increased acH3K9 expression in the core. Thus, adolescent cocaine experience interacts with genetic background to elicit different behavioral profiles relevant to addiction in adulthood, with concurrent modifications in the epigenetic histone profiles in the nucleus accumbens that associate with cocaine sensitization and with metaplasticity.

**SIGNIFICANCE STATEMENT:** Adolescence is a critical period when brain developmental trajectories continue to be established and when most illicit drug use is initiated in humans. Notably, however, only a minority of adolescent drug users go on to develop substance use disorders in adulthood. We hypothesize that individual differences in genetic vulnerability, temperamental phenotype and adolescent drug history intersect to modify life-long drug abuse liability, with epigenetic modification playing an important role in this interplay. We tested this hypothesis in a genetic animal model of temperament. Our results show that adolescent cocaine experience induces psychomotor sensitization in vulnerable individuals, and that a dual exposure to cocaine (both in adolescence and adulthood) limits further sensitization in these individuals, an example of metaplasticity. By contrast, individuals who are typically resilient become more responsive to cocaine when exposed to this drug during adolescence and are then re-exposed as adults. Differences in behavioral response to cocaine were associated with unique epigenetic changes in the nucleus accumbens core. These findings shed light on the interplay of genes, environment and adolescent exposure to drugs in susceptibility to substance use disorders.

## Introduction

Substance use disorder (SUD) is a complex, pernicious and pervasive condition negatively impacting millions around the world and costing hundreds of billions of dollars in the United States alone (Volkow et al., 2016; NSDUH, 2020). However, despite years of research, widely effective treatments remain elusive (Lipari et al., 2012; http://www.samhsa.gov/). One reason may be the confounding contribution of individual differences. SUD is highly heritable (30-70%; Ducci and Goldman, 2012), and interrelated with other psychiatric diseases, often sharing common neurobiology (Goldman et al., 2005; Kreek et al., 2005; Hawkins, 2009). Twin studies suggest SUD and externalizing disorders, including novelty seeking, have shared genetic influence (Hicks et al., 2004; Hettema et al., 2005), while early-onset depressive or internalizing disorders appear to be inherent predisposing factors, often co-morbid in persons with SUD and predictive drug dependence problems later in adulthood (Crews et al., 2007; Sihvola et al., 2008; Chen et al., 2009). This suggests that temperamental tendencies likely contribute to drug use and misuse likely through multiple paths - e.g., novelty seeking in some cases, and self-medication for negative affect in others. Moreover, environmental factors, such as drug availability, childhood adversity, socioeconomic distress, and the impact of chronic drug experience are strongly associated with addiction outcome. Understanding how gene x environment interactions contribute to addiction liability is therefore an important area of investigation that will help improve treatment options for individuals with SUD as well as informing prevention strategies.

Most individuals with SUD initiate drug use before the age of 18 and then quickly escalate use, resulting in a life-long problem by adulthood (Warner, 1995; Degenhardt et al., 2007; Burstein, 2012). Younger age of onset often predicts greater risk of SUD later in life (Chen et al., 2009; McCabe et al., 2007; Palmer et al., 2009) and drug abuse during adolescence, a sensitive period when these environmental and stress-related effects are most formative, seems to be a consistent factor in developing long-term addiction liability (Dickinson and Forsyth, 1994). However, despite these relationships, only a small minority of individuals who use illicit drugs develop SUD. Therefore, it is likely that a critical combination of individual differences (i.e., inherent predisposing and environmental risk factors) work in concert to establish lifelong addiction vulnerability (Rao and Chen, 2008).

The field of neuroepigenetics provides biological evidence that genes are accompanied by heritable epigenomic profiles (e.g., chromatin structure) that can become altered by life experiences, including chronic drug use (Nestler, 2014; Pierce et al., 2018). Thus, under the right circumstances even inborn genetic transcription profiles can become modified by environmental experience, altering inherent affective phenotypes (McGowan and Szyf, 2010; Kanherkar et al., 2014; Kundakovic and Champagne, 2015; McEwen, 2018). In agreement, several groups have shown that a history of drug experience can induce epigenetic modifications that influence gene expression and contribute to addiction vulnerability both within a lifetime (LaPlant and Nestler, 2011; Feng and Nestler, 2013; Maze et al., 2011) and across generations (Vassoler et al., 2014; Le et al., 2017; Vassoler and Sadri-Vakili, 2014). While epigenetic modifications occur throughout the brain, changes in particular regions within the striatum (e.g., the nucleus accumbens, NAc, core) have been shown to directly influence dopamine-rich mesolimbic medium spiny neurons, which play a role in addiction-related behaviors (Kennedy et al., 2013). For example, selective histone modification on Histone 3 Lysine 9 (H3K9) in the NAc core, manipulated via engineered zinc finger proteins targeting Cdk5 gene expression, mediates cocaine conditioned place preference and psychomotor sensitization in mice (Heller et al., 2016).

Adolescence is a critical period where chromatin structure is being modified by hormonal and other factors, and extensive drug use during adolescence is likely to have a strong effect on chromatin structure, possibly impacting the neurobiological substrates that contribute to addiction-related phenotypes. However, the interplay between genetic background and the epigenetic response to chronic drug exposure, especially during adolescence, remains understudied. Here, we used our bred High-responder (bHR) and Low-responder (bLR) rat model to investigate whether genetic predisposition intersects with adolescent cocaine experience to alter addiction-related behaviors and epigenetic profiles in adulthood.

Our two lines of rats are bred based on their locomotor response to a novel environment, but concurrently differ on a number or addiction-related measures, including psychomotor sensitization, acquisition of drug-taking, response to drug cues, and dopamine receptor 1 (D1) and 2 (D2) RNA profiles in the NAc (Davis et al., 2008; Flagel et al., 2010; Clinton et al., 2011; Flagel et al., 2014). Now bred across multiple generations, these rats reliably exhibit distinct and extreme novelty-seeking and addiction-related phenotypes that are predicted with 99% certainty (Flagel et al., 2014). This a priori knowledge allowed us to test whether adolescent cocaine history augments or redirects their respective inborn phenotypes and track changes to their epigenetic profiles. We measured psychomotor sensitization and profiled two opposing H3K9 modifications (acetylation and tri-methylation) previously implicated in addiction-related behaviors across four striatal sub-regions of bHR and bLR rats following exposure to cocaine (or saline): 1) in adolescence only, 2) in adulthood only, or 3) in both adolescence and adulthood. While most other studies examined epigenetic changes in bulk, at the regional level, we sought to examine the changes at a more cellular level within the striatum. Thus, we relied on a combination of immunohistochemistry and unbiased stereology to more precisely visualize the epigenomic changes associated with exposure to cocaine.

## Material and methods

### Animals

A total of 150 male Sprague-Dawley rats (70 bHR and 80 bLR) from generation F43 were used in this study. For unknown reasons, one bLR rat died prior to initial locomotor testing and his cage-mate was subsequently excluded. Rats were housed in pairs in standard controlled environmental conditions (22 °C, 70% humidity, and 12-h light/dark cycle, lights on at 06:00 h) with food and water available *ad libitum*. All animal procedures were performed in accordance with the University of Michigan animal care committee’s regulations and followed the Guide for the Care and Use of Laboratory Animals: Eight Edition, (Committee for the Update of the Guide for the Care and Use of Laboratory Animals, 2011).

### Basal Locomotor Response to Novelty

All rats were tested for locomotor response to a novel environment, the phenotype for which bHR and bLR rats were bred, during early life (P25) between 9:30 AM and 12:30 PM in a different room from where the rats were housed. Rats were placed into clear acrylic 43 × 21.5 × 25.5 cm tall cages equipped with infrared photocell emitters mounted 2.3 and 6.5 cm above a grid floor placed on top of standard bedding. A customized computer with a locomotion-testing rig and motion-recording software created in-house at the University of Michigan was used to measure lateral and rearing activity. Each locomotor test session lasted 60 min and the data were collected and recorded in 5-min bins. Locomotion scores for each rat were calculated as the sum of horizontal and rearing movements for the entire session. Recorded locomotor scores were used to counterbalance all subsequent testing conditions within bHR or bLR phenotypes (see Supplementary Fig S1 and Table S1).

### Drug Treatments and Psychomotor Activating Effects of Cocaine

A total of 60 bHR and 78 bLR rats were intraperitoneally administered either cocaine (15 mg/kg) or saline once daily for 7 consecutive days during adolescence (P33-39) (Fig 1). While a cohort of rats was sacrificed 24 h after treatment (P40) (20 bHR and 38 bLR; see Fig. 1), the other cohort was exposed to the same regimen of either saline or cocaine administration in adulthood (P76-84), providing 8 experimental groups (see n/group in Fig. 1). Another group of 10 bHR rats was only exposed to cocaine in adulthood (no drug treatment during adolescence; ND-Cocaine; Fig. 1). This selected cocaine regimen was previously shown to induce psychomotor sensitization in adult outbred (Gosnell et al., 2005) and bHR rats (García-Fuster et al., 2010), and to induce long-term behavioral changes (e.g., Garcia-Cabrerizo et al., 2015; García-Fuster et al., 2017; García-Cabrerizo and García-Fuster, 2019) when given at this time-window during early-mid adolescence.

**Fig. 1.**
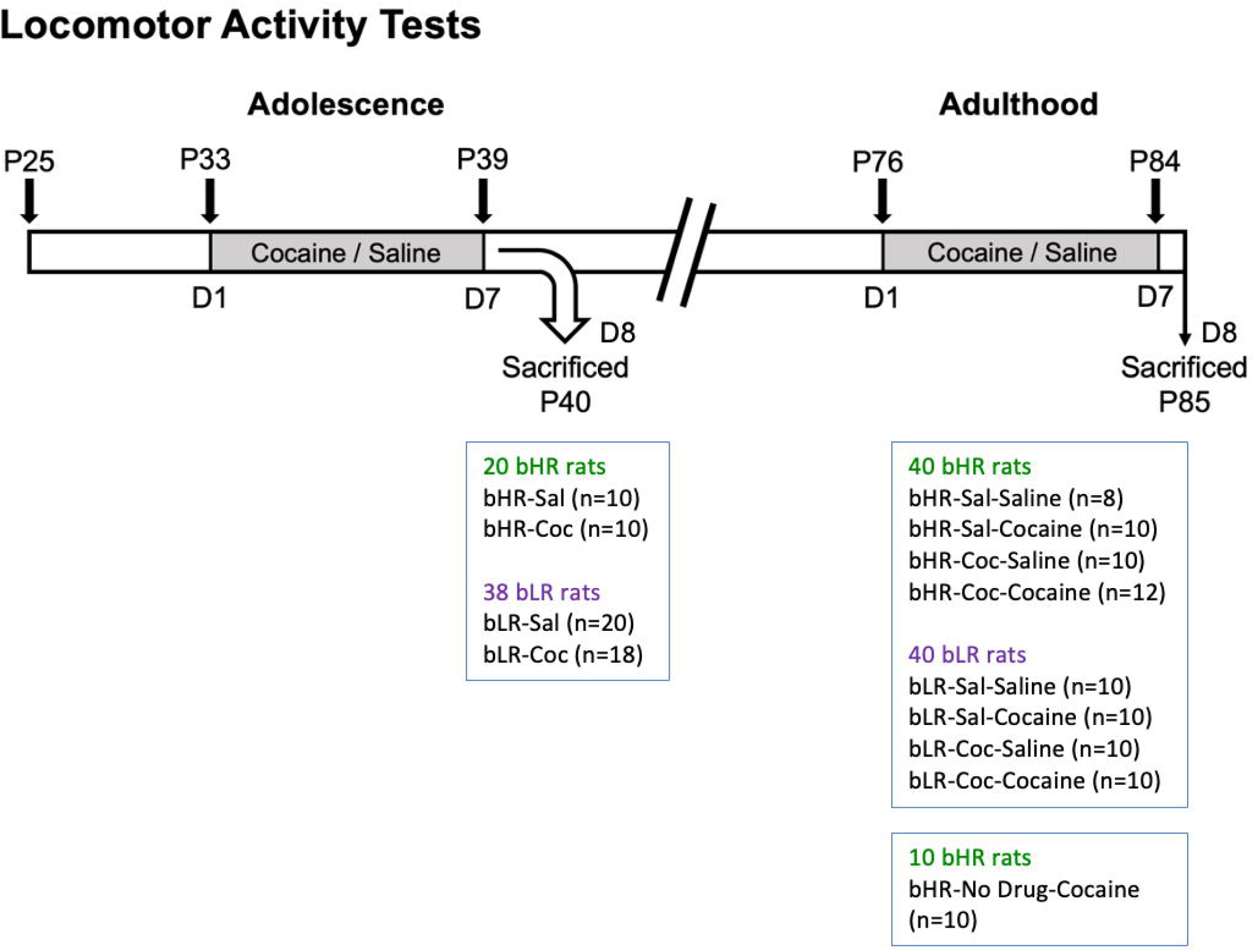
Drug Treatments and Psychomotor Activating Effects of Cocaine. Locomotor Activity Tests. P#: post-natal day; Gray: drug treatment period. bHR and bLR rats were initially tested for locomotor exploration of a novel environment (P25, Supplementary Fig. S1). bHRs and bLRs were then assigned to different treatment conditions where either saline or cocaine was administered in adolescence (P33-39), adulthood (P76-84), or both. A subset of these bHRs and bLRs were sacrificed 24 h (D8, P40) following the adolescent cocaine sensitization regimen to assess epigenetic changes at this timepoint. All other conditions were administered the second regimen of either saline or cocaine later in adulthood (P76-P84) and subsequently sacrificed (P85).

Psychomotor sensitization tests were performed during adolescence, adulthood, or both. On the first day (D1) of each sensitization regimen (P33 or P76), rats were transferred to a dedicated testing room where they were placed into custom-made activity chambers (García-Fuster et al., 2017) and were allowed to habituate for 2 h before receiving an injection of either cocaine (15 mg/kg, i.p.) or saline (0.9% NaCl, 1 ml/kg, i.p.). This was immediately followed by 1 h of locomotor testing before being returned to their home cages. For the next 5 days (P34-38, and/or P77-83), rats were injected once daily with 15 mg/kg of cocaine or saline in an alternate room at the same time of day as the first test. Rats were placed back into their homecages immediately following injections and returned to their colony room. On D7 of testing (P39 and/or P84), rats were again returned to the activity chambers where they were tested on D1 and allowed to habituate for 2 h before receiving an injection of either cocaine or saline followed by 1 h of behavioral scoring.

### Test Equipment and Analysis of Signs of Psychomotor Sensitization

As previously described (García-Fuster et al., 2017) locomotor activity test chambers were made of expanded PVC (33.02 × 68.58 × 60.96 cm tall) with stainless steel woven-wire cloth grid floors (30.48 × 60.96, 7.62 × 7.62 cm squares), placed above a metal catch tray. Animal behavior was digitally video recorded using IC Capture 2.2 software (www.theimagingsource.com) and USB 2.0 monochrome industrial video cameras (The Imaging Source LLC, Charlotte, NC) installed directly above each activity chamber. Recordings captured behavior during a 2-hour habituation period and for 1 hour following drug administration. These video files were transferred to an offline computer server for subsequent behavior analysis using TopScan 2.0 software (Clever Sys, Inc.; Reston, VA, USA, www.cleversysinc.com). Several measurements were used to ascertain signs of psychomotor sensitization as described by Flagel and Robinson (2007). Locomotor activity was measured as the distance travelled in mm from one end of the chamber to the other based on user-defined areas of the activity chamber, and the velocity (mm/s). The number of darting bouts, which is defined as an uninterrupted instance of fast locomotion (≥ 95 mm/s as the predetermined velocity) was also measured. Finally, stereotypy was measured by TopScan using the frequency of head-waving movements: the total number of lateral head-waving movements divided by the time spent in place (no./s).

### Tissue Collection

Rats were killed by rapid decapitation using a guillotine 24 h following the last day of cocaine or saline administration during adolescence (D8: P40) or in adulthood (D8: P85; see Fig. 1). The extracted brains were quickly frozen in a −30 °C isopentane solution and stored at −80 °C until further processing. For each rat, 40 ¼m coronal sections were cryostat cut through the rostrocaudal extent from cortex to the brainstem and slide-mounted (Superfrost Plus, VWR). Mounted sections were then stored at −80 °C for subsequent immunohistochemistry (IHC) and/or non-radioactive in situ hybridization assays.

### Histone Modification Quantification Visualization

Tri-methyl and acetylated H3K9 epigenetic marks (H3K9me3 and acH3K respectively) were systematically quantified following diaminobendazine (DAB) immunohistochemistry for regional quantification in stritatal regions (Fig. 2). Immunohistochemistry experiments followed a standardized protocol (García-Fuster et al., 2010) where tissue was post-fixed in 4% paraformaldehyde and incubated at 90 °C for 50 minutes in 0.1 M sodium citrate (Fisher, #S379-3, pH 6.0) for epitope retrieval. Sections were then rinsed with Tris-buffered saline (TBS), washed in 0.3% peroxide, blocked with bovine serum albumin containing 1% goat serum and 0.05% Triton X-100, and incubated overnight with either monoclonal rabbit anti-H3K9me3 (1:2000; Abcam, ab8988, USA) or polyclonal anti-acH3K9 (1:5000; Cell Signaling, #9649, USA). After PBS washes, sections were incubated in biotinylated anti-rabbit secondary antibody 1:500 (Vectors Laboratories, #BA-1000) followed by Avidin/Biotin complex amplification (Vectastain Elite ABC kit; Vectors Laboratories, #PK-6100) and DAB (Sigma Life Science, #5905) mixed with 3% hydrogen peroxide for signal amplification. Finally, all slides underwent dehydration through graded alcohols, xylene immersion, and were subsequently coverslipped using Permount® mounting medium (Fisher, #SP15).

**Fig. 2.**
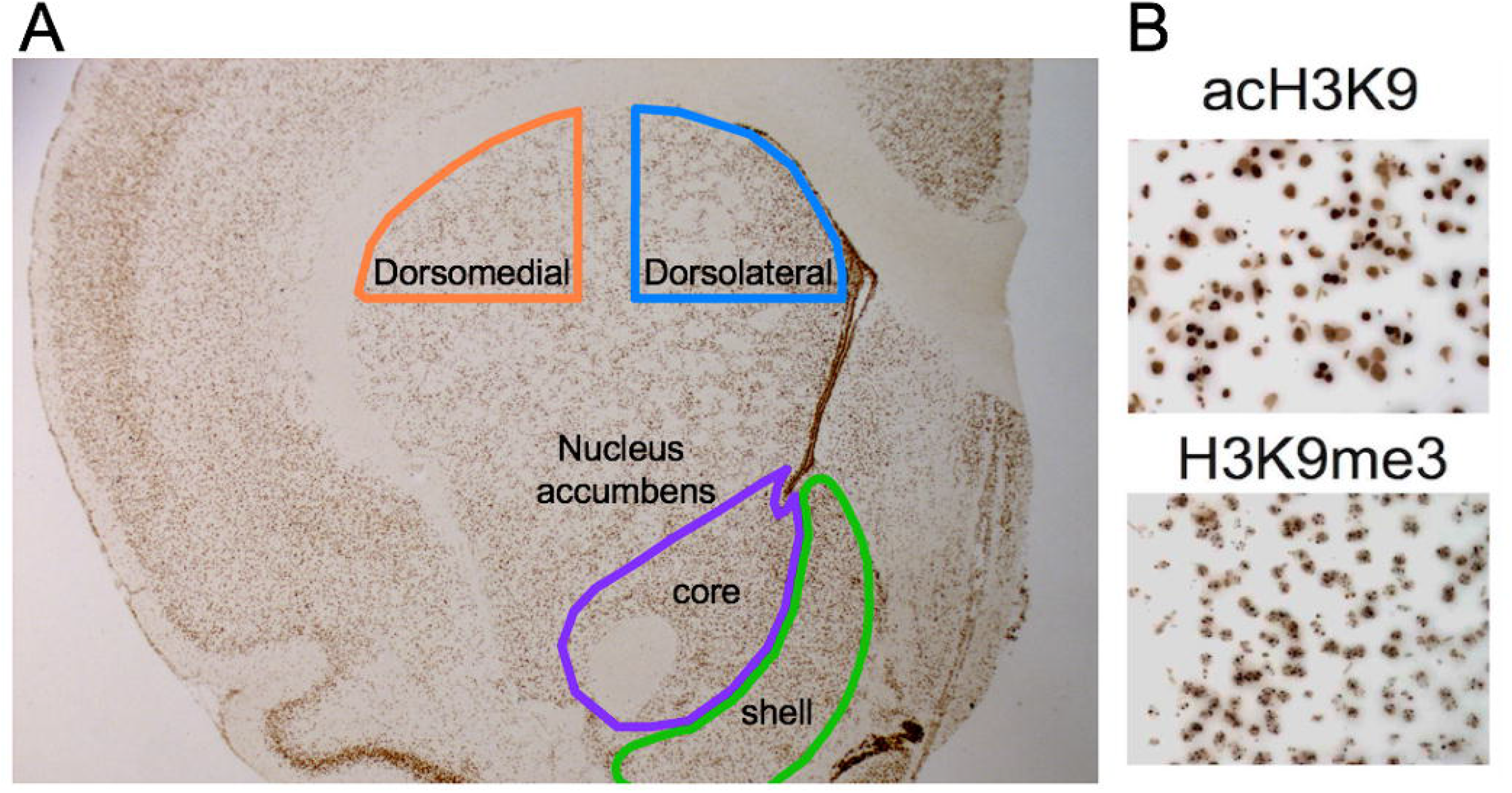
Histone Modification Quantification Visualization. A modified combination of immunohistochemistry and unbiased stereology was used to quantify acH3K9 and H3K9me3 histone modifications within defined anatomical sub-regions of the striatum: dorsomedial (dmSTR), dorsal lateral (dlSTR), nucleus accumbens (NAc) core and shell. Each area was outlined with hand-drawn contours based on Paxinos and Watson brain atlas (2012) (A), and then quantified using unbiased stereology (see methods). Antibody concentrations for acH3K9 and H3K9me3 were optimized using a titration curve and visualized using diaminobenzidine (DAB) for easy counting (B).

### Unbiased Stereology

Unbiased stereology was performed using a Zeiss Axioimager M2 motorized fluorescent microscope at 40x or 63x oil. DAB immunohistochemistry-processed slides representing every animal were randomized and coded prior to analysis. Trained personnel blind to the treatment conditions performed all of the stereology. Systematic Random Sampling (SRS) was performed on each section using Stereoinvestigator software (MBF) with frames (60 × 60 µm) on a counting grid (150 × 150 µm). Optical dissector height was 20 µm with 2 µm upper and lower guard zones to avoid quantification of partial cells and cutting artifacts. All counting protocols used a Gundersen coefficient (m=1) of <0.1 and near equal values between treatment conditions (CE= ∼0.04) to ensure count accuracy and consistency (Gundersen et al., 1999). Some brains did not stain well enough for optimal stereological quantification, but attrition was random, small, and no condition had fewer than 8 rats for analysis of epigenetic modifications.

### Statistical Analysis

All graphical representations of the data and the corresponding statistical analysis were performed with GraphPad Prism, Version 9 (GraphPad Software, San Diego, CA). Results are expressed as mean values ± standard error of the mean (SEM) and individual symbols for each rat are shown within bar graphs. Given their baseline differences in locomotor activity between bHR and bLR rats (Fig. S1), the statistical comparisons evaluating the psychomotor effects of cocaine were performed in bHR and bLR separately. In particular, following the adolescent treatment, two-way repeated measures (RM) ANOVAs or mixed-effects models (if there were any missing values) were used with the following independent variables: Adolescent Treatment and Day of testing (D1 vs. D7), and followed by Sidak’s multiple comparison tests when appropriate. Following the adult treatment, three-way RM ANOVAs were performed, which included 3 independent variables (Adolescent Treatment, Adult Treatment, Day of testing; see Supplementary Table S1). When evaluating the changes in epigenetic markers (H3K9me3 and acH3K9) in the NAc (core and shell), Student *t*-tests were performed to assess the effects of adolescent cocaine (D8, 24 h post-adolescent treatment), while two-way ANOVAs were utilized (independent variables: Adolescent Treatment and Adult Treatment) following the adult treatment (D8, 24 h post-adult treatment). Sidak’s multiple comparisons tests were used for *post-hoc* comparisons when appropriate. For statistical analysis during adulthood 4 experimental groups (Sal-Saline, Sal-Cocaine, Coc-Saline, Coc-Cocaine) were included; however, graphs were split accordingly to history of cocaine for simplicity and visualization. The level of significance was set at *p*≤0.05.

## Results

### Adolescent Cocaine Differentially Impacts Psychomotor Sensitization in Divergent Affective Phenotypes

#### Effects during adolescence

During adolescence, both acute (D1) and chronic (D7) cocaine treatment regimens elicited psychomotor activation in bHR (Fig. 3A-B) and bLR (Fig. 3C-D) rats (statistical results are reported in Supplementary Table S1). For each phenotype there was a significant main effect of Treatment, supported by increased locomotor activity (i.e., distance travelled, Fig. 3A and 3C; locomotor velocity, darting bouts, Fig. S2) and head-waving frequency (Fig. 3B and 3D) on D1 and D7. However, only for bHR rats was there a significant effect of Day (D7 vs. D1) for locomotor activity (see Supplementary Table S1). Thus, following repeated drug exposure in adolescence only bHRs show evidence of psychomotor sensitization (Fig. 3A).

**Fig. 3.**
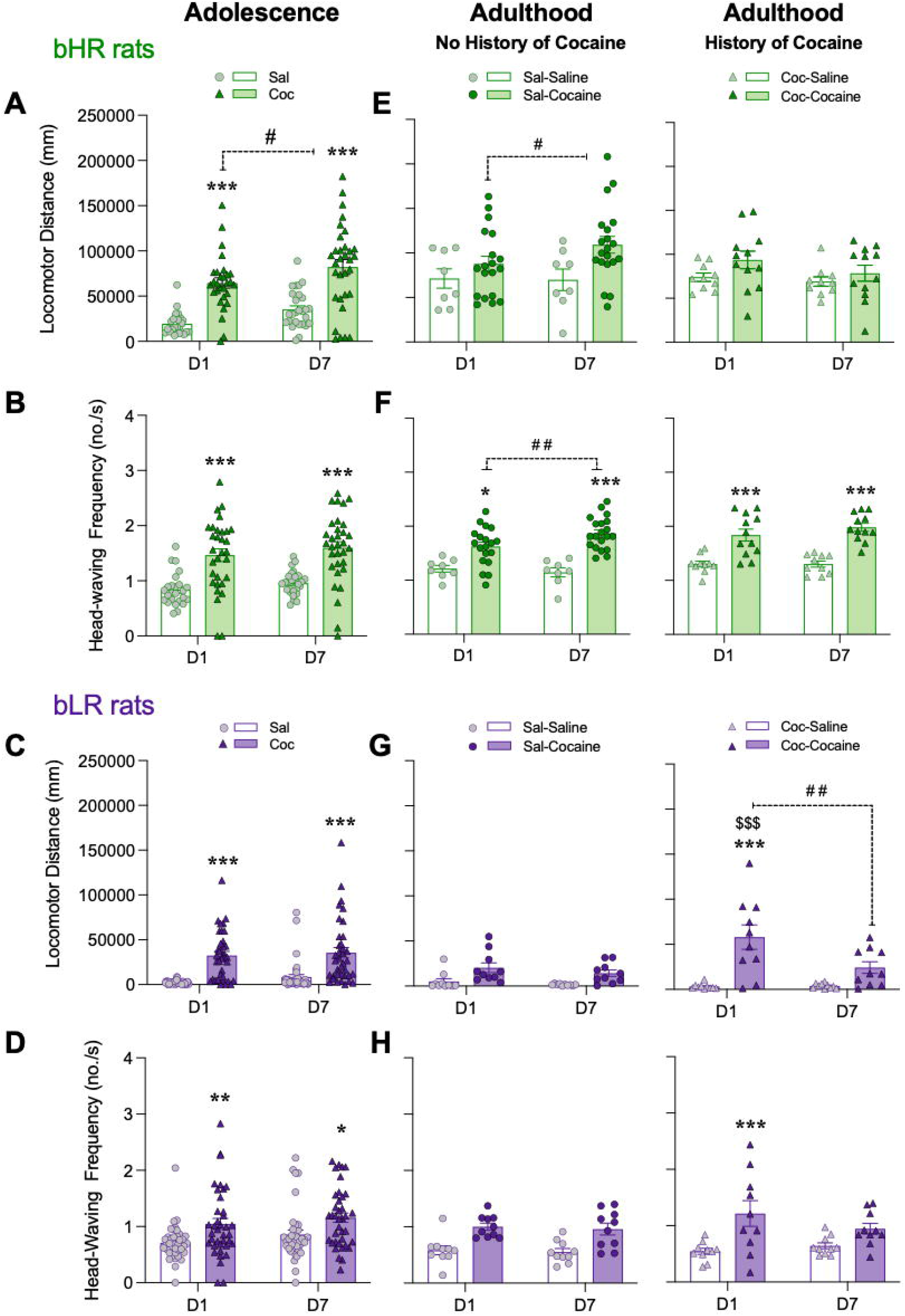
Adolescent Cocaine Differentially Impacts Psychomotor Sensitization in Divergent Affective Phenotypes. Acute (D1) and chronic (D7) psychomotor activating effects of cocaine administration in adolescent (panels A-D) and/or in adulthood (with no or with a prior history of adolescent cocaine; panels E-H) in bHR and bLR rats. Data represents the mean values ± SEM of the locomotor distance travelled in mm (panels A, C, E and G) and the frequency of head-waving (panels B, D, F and H). Individual values are shown for each rat (symbols). Two- or three-way RM ANOVAs were used for statistical analysis (see Supplementary Table S1 for more details). \**p* < 0.05, \*\**p* < 0.01 and \*\*\**p* < 0.001 when comparing cocaine-treated rats with the corresponding saline-treated group (main Effect of Treatment). ##*p* < 0.01 and #*p* < 0.05 when comparing D7 vs. D1 responses for the same treatment group (main Effect of Day). $$$*p* < 0.001 when comparing the response on D7 of the adolescent treatment vs. the one following an acute challenge dose in adulthood (D1 for Coc-Cocaine group).

#### Effects during adulthood

In bHRs, the psychomotor effects of adult cocaine exposure where only observed for locomotor distance (Fig. 3E) and head-waving frequency (Fig. 3F) (see the overall main effects of Treatment and other statistical results reported in Table S1). In bHR rats there was a significant Day x Adult Treatment interaction when evaluating head-waving frequency (see Table S1), and Sidak’s multiple comparisons test revealed an increase in head-waving frequency on D7 as compared to D1 for rats treated with cocaine (see Fig. 3F).

However, bHRs exposed to adult cocaine and with a prior history of adolescent cocaine showed similar locomotor activity levels (i.e., locomotor distance and velocity, darting bouts) and head-waving frequency movements on D1 and D7 in adulthood, and as compared to rats that only received cocaine in adulthood (Fig. 3E-F and Supplementary Fig. S2). Thus, bHR rats that were exposed to cocaine in adolescence do not exhibit further sensitization in adulthood.

In bLR rats with a history of cocaine exposure during adolescence, psychomotor effects of adult cocaine treatment were observed for all measurements (i.e., locomotor distance and velocity, darting bouts, head frequency; see main effects of Adult Treatment on Supplementary Table S1 and Figs. 3G-H and S2). These rats showed greater locomotor activity and head-waving frequency in response to a single cocaine injection (D1) relative to those treated with a repeated cocaine treatment during adolescence or an acute dose of cocaine in adulthood (Fig. 3G-H). Notably, the increase in locomotor activity on D1 was followed by a reduction on D7 (Fig. 3G), suggesting a lack in the classic pattern of adult sensitization (increased activity from D7 vs. D1).

### Adolescent Cocaine Differentially Impacts Epigenetic Expression in the Nucleus Accumbens Core of Divergent Affective Phenotypes

Prior to evaluating the impact of drug treatments in adolescence and/or adulthood, we ascertained potential baseline differences in the expression of H3K9me3 and acH3K9 with age (adolescence P40 vs. adulthood P85) in saline-treated rats. There were no significant differences between bHR and bLR rats at baseline in either histone marker in the NAc core or NAc shell (Supplementary Fig. S3).

#### Effects during adolescence

Chronic adolescent cocaine treatment reduced H3K9me3 in the NAc core of bHR rats (−42476 ± 15169, \**p*=0.0150 vs. Sal treated rats; Fig. 4A), but not bLR rats (Fig. 4B), when measured 24 h after the last dose (D8) (see Supplementary Table S1). In contrast, there was an increase in acH3K9 in the NAc core (+24757 ± 12914, \**p*=0.0421 vs. Sal treated rats) in bHR rats treated with cocaine (Fig. 4E), but not bLR rats (Fig. 4F). There were no significant effects of cocaine treatment on either H3K9me3 (Fig. 3C-D) or acH3K9 (Fig. 3G-H) in the NAc shell in bHR or bLR rats 24 h after the last treatment.

**Fig. 4.**
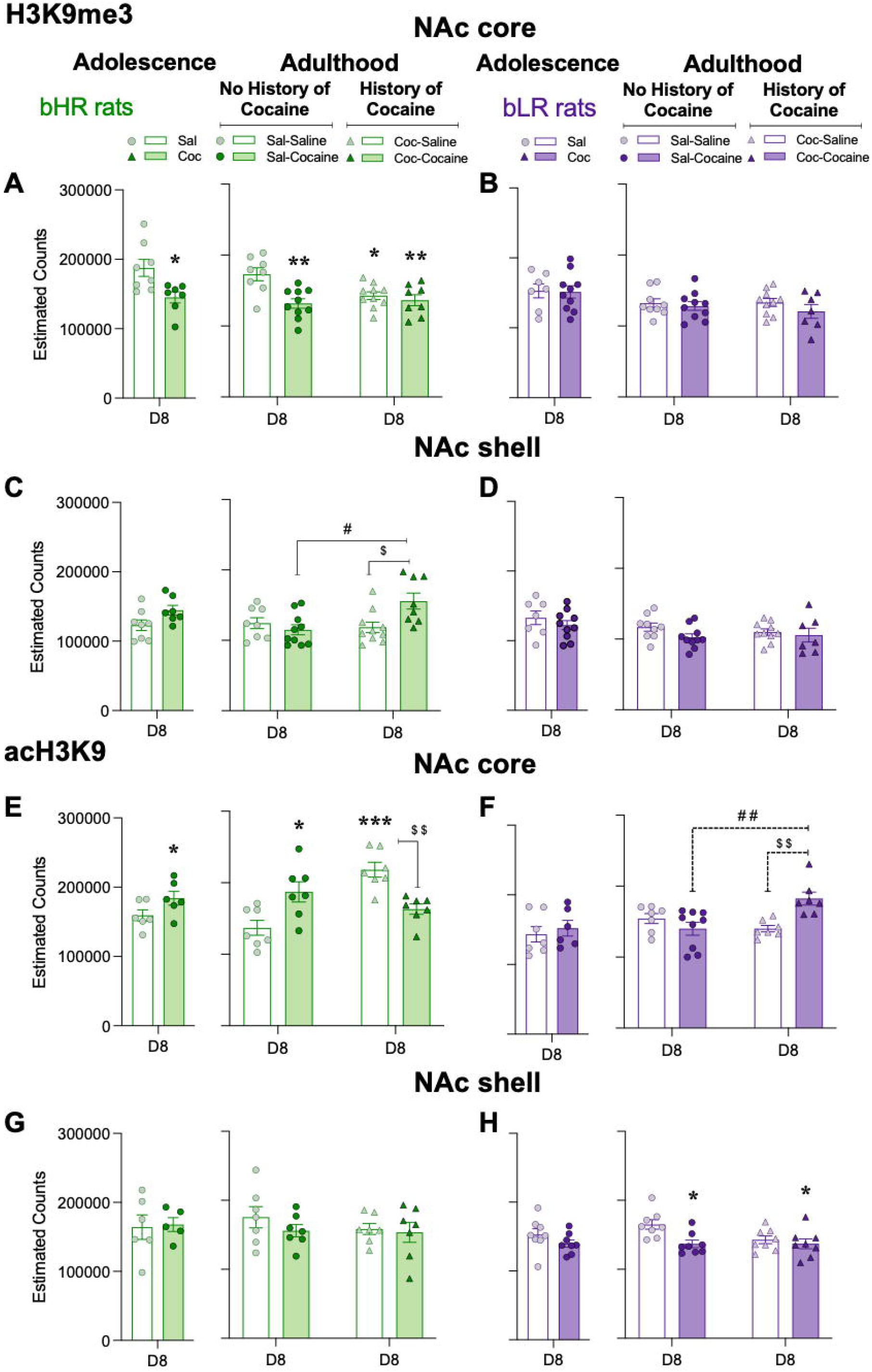
Adolescent Cocaine Differentially Impacts Epigenetic Expression in the Nucleus Accumbens Core of Divergent Affective Phenotypes. Repressive H3K9me3 (panels A-D) and permissive acH3K9 (panels E-H) cell counts in the NAc core and shell of bHRs and bLRs treated with adolescent and/or adult cocaine. Data represents the mean values ± SEM of the estimated cell counts of each marker as measured 24 h post-adolescent or adult treatment (D8). Individual values are shown for each rat (symbols). Student *t*-tests were performed to assess the effects of adolescent cocaine, while two-way ANOVAs were utilized following the adult treatment (see Supplementary Table S1 for more details). \**p* < 0.05 and \*\**p* < 0.01 when comparing cocaine-treated rats with the corresponding saline-treated group (main Effect of Treatment). ##*p* < 0.01 and #*p* < 0.05 when comparing rats with and without a prior history of adolescent cocaine (Coc-Cocaine vs. Sal-Cocaine). $$*p* < 0.01 and $*p* < 0.05 when comparing Coc-Cocaine vs. Coc-Saline groups.

#### Effects during adulthood

In adulthood, H3K9me3 was again decreased in the NAc exclusively in bHR rats (Fig. 4A). Specifically, two-way ANOVAs revealed Adolescent x Adult Treatment interactions in both the NAc core and NAc shell (see Supplementary Table S1). In the NAc core (see Fig. 4A), Sidak’s multiple comparisons tests showed that adult cocaine administration reduced the expression of H3K9me3 in bHR rats with (Coc-Saline: −30162 ± 10092, \**p*=0.0317 vs. Sal-Saline) and without (Sal-Cocaine: −41123 ± 10092, \*\**p*=0.0017 vs. Sal-Saline) a history of adolescent cocaine. The combination of cocaine exposure during both adolescence and adulthood resulted in decreased expression of H3K9me3 (Coc-Cocaine −36574 ± 10638, \*\**p*=0.0098 vs. Sal-Saline), and with a similar magnitude of change between those groups that received cocaine during adolescence and those that received cocaine in adulthood (Fig. 4A). In the NAc shell (Fig. 4C), the combination of adolescent and adult cocaine treatment (Coc-Cocaine) resulted in higher levels of H3K9me3 in bHR rats, compared to those that received cocaine only during adolescence (+37223 ± 11252, $*p*=0.0227 vs. Coc-Saline) or in adulthood (+40755 ± 11935, #*p*=0.0105 vs. Sal-Cocaine).

For acH3K9 in the NAc, two-way ANOVAs revealed Adolescent x Adult Treatment interactions in the NAc core for both bHR (Fig. 4E) and bLR (Fig. 4F) rats (see Supplementary Table S1 for statistical analysis). In bHR rats, a single regimen of cocaine exposure (either in adolescence or adulthood) increased acH3K9 (Sal-Cocaine: +50299 ± 14234, \**p*=0.0179; Coc-Saline: +81239 ± 15234, \*\*\**p*=0.0001) relative to saline-treated (Sal-Saline) rats (Fig. 4E). However, acH3K9 was not affected in bHR rats that received a combination of adolescent and adult cocaine treatment (Coc-Cocaine: −55097 ± 15234, ##*p*=0.0082 vs. rats that just receiving adolescent cocaine, Coc-Saline; Fig. 4E). In bLR rats, the combination of adolescent and adult cocaine (Coc-Cocaine) treatment resulted in higher expression of acH3K9 in the NAc core when compared to those that only received treatment during adolescence (+42325 ± 11627, $$*p*=0.0071 vs. Coc-Saline) or in adulthood (Sal-Cocaine: +42494 ± 10962, ##*p*=0.0039) cocaine treatments (Fig. 4F). In the NAc shell, acH3K9 was decreased in bLR rats following cocaine treatment in adulthood (Fig. 4H).

To further examine the long-term impact of adolescent cocaine exposure on brain regions involved in psychomotor sensitization, we also quantified the number of cells expressing acH3K9 in the dorsomedial (dmSTR) and dorsolateral (dlSTR) regions of the striatum in adult bHR and bLR rats (Fig. S4). There were no significant changes induced by cocaine exposure (adolescent, adult or both) in these subregions in either bHR or bLR rats (Fig. S4; Table S1).

## Discussion

In the present study, we analyzed the interplay of genetic predisposition and exposure to cocaine during adolescence in shaping behavioral responsiveness to cocaine during adulthood. We asked whether the unique behavioral phenotypes that are shaped by these gene x environment interactions are associated with distinct epigenetic profiles. In particular, we studied whether exposure to chronic cocaine alters the balance of a repressive and a permissive modified histone differentially, as a function of genetic background.

Our <u>behavioral findings</u> show the following: a) bHR rats, which are vulnerable to psychostimulant use and exhibit a number of addiction-related behaviors (Flagel et al., 2010, 2014, 2016) are prone to sensitization in both adolescence and adulthood across several measures of locomotor behavior; b) In bHRs, previous exposure to cocaine sensitization during adolescence abrogates the development of sensitization in adulthood, indicating the existence of limits on drug-induced neuroplastic changes -- a phenomenon known as metaplasticity; c) Adult bLRs, which are comparatively resilient to psychostimulant use, are only mildly reactive to either acute or chronic cocaine. By contrast, adolescent bLRs show greater locomotor activation in response to acute and chronic cocaine, although they are still less responsive than bHRs. However, at neither age do bLRs show cocaine sensitization in the sense of an increased locomotor response on D7 relative to D1; d) In bLRs, a previous exposure to cocaine during adolescence produces an unusual pattern of reactivity in adulthood. The initial cocaine challenge in adulthood elicits a pronounced locomotor response in comparison to bLRs with no history of cocaine. This is evidence of sensitization triggered by adolescent exposure that becomes manifest on the initial cocaine challenge in adulthood. However, following 7 days of exposure to cocaine in adulthood, this enhanced initial response appears resistant to further sensitization on most measures of locomotion, and shows a significant reduction in locomotor distance traveled. Thus, the adolescence-induced sensitization to cocaine in adult bLRs appears to be transient. Overall, cocaine exposure during adolescence caused enduring effects into adulthood with distinct sensitization phenotypes as a function of genetic predisposition.

Our <u>epigenetic findings</u> show the following: a) In bHRs, sensitization was associated with a decrease in the repressive H3K9me3 and an increase in the permissive acH3K9 in the NAc core, a combination expected to lead to an increase in net transcriptional activation. These sensitization-induced changes appear to be persistent (i.e., bHR with a history of adolescent cocaine who receive saline as adults have a profile resembling the adolescent sensitization profile, with decreased H3K9me3 and increased acH3K9); b) In bHRs with dual exposure to cocaine in adolescence and adulthood, the modified histones showed either no additional changes (H3K9me3) or a reversal (acH3K9), consistent with metaplasticity; c) In bLRs, there are no decreases in H3K9me3 in the NAc core in any of the experimental groups; d) In the bLRs, there is an increase in the permissive acH3K9 in the NAc core only in adults with a history of cocaine exposure during adolescence. It is notable that this group showed evidence of transient sensitization to cocaine.

Overall, sensitization in its typical manifestation seen in bHRs is associated with a decrease in the repressive H3K9me3 and increase in a the permissive acH3K9. Limits on their changes are associated with limits to behavioral sensitization (see Fig. 5). By contrast, an increase in acH3K9 alone, while associated with an increased responsiveness to an initial cocaine challenge in bLRs is not sufficient to support the expression of sustained sensitization. These findings, and additional behavioral and epigenetic observations, are further discussed below.

**Fig. 5.**
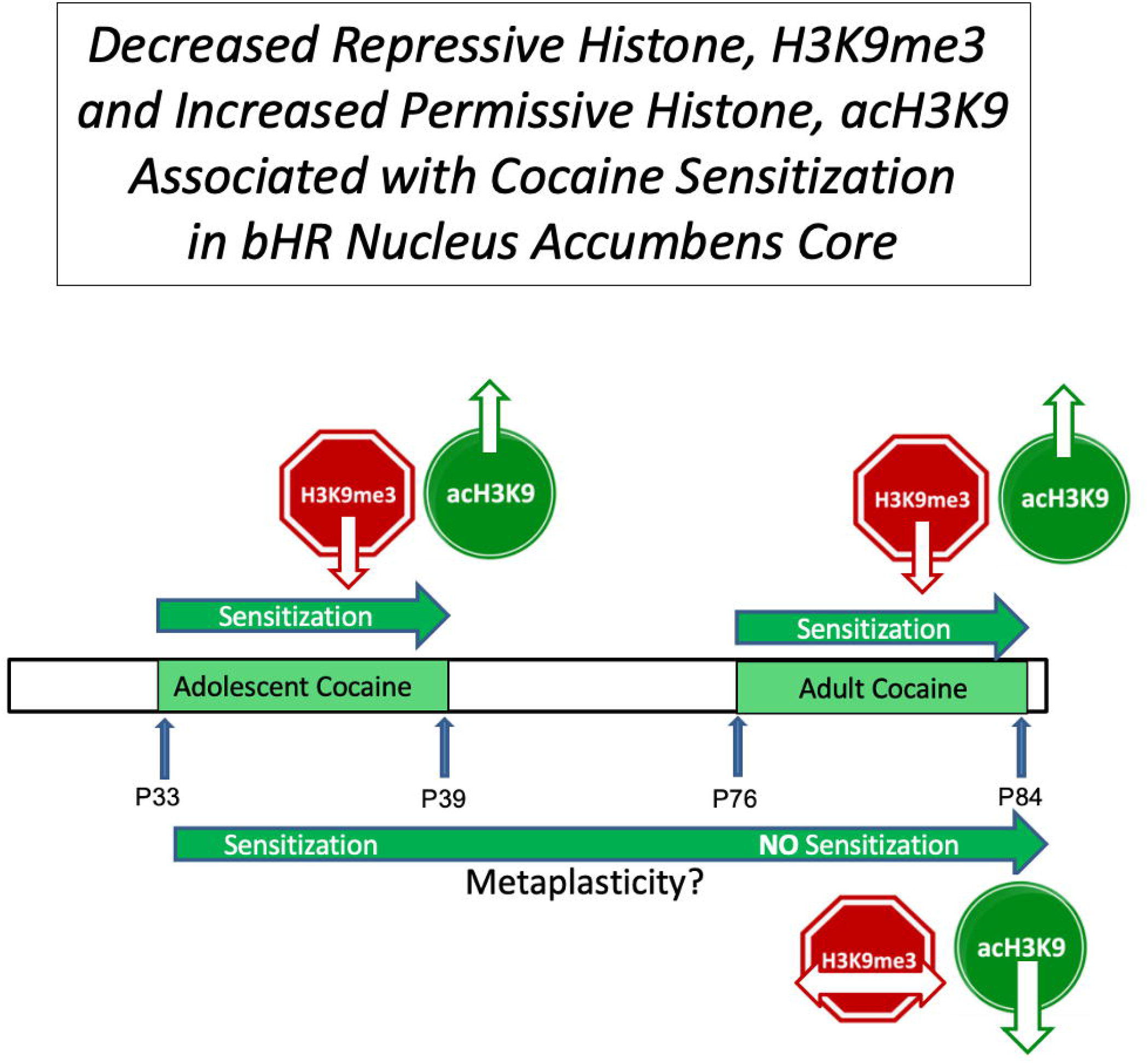
Decreased Repressive Histone, H3K9me3 and Increased Permissive Histone, acH3K9 Associated with Cocaine Sensitization in bHR Nucleus Accumbens Core. Illustration that summarizes the coordinate changes in the repressive H3K9me3 (red) and permissive acH3K9 (green) histones in relation to sensitization and metaplasticity for bHR rats. Downward arrow indicates a decrease, upward arrow an increase, and horizontal arrow no change. bHR rats with a history of cocaine exposure in adolescence show no change in H3K9me3 in adulthood and no evidence of behavioral sensitization, consistent with the notion of metaplasticity.

### The Impact of Adolescent and Adult Exposure to Cocaine on Behavioral Sensitization

bHR rats are largely considered more addiction liable as they inherently exhibit greater risk-taking and exploratory behaviors, more readily acquire self-administration of cocaine, show greater spontaneous dopamine release, higher D2 mRNA expression in the dorsal striatum, and greater locomotor responsiveness to psychostimulants, which may facilitate initiation of drug seeking (Flagel et al., 2010; Flagel et al., 2014; Flagel et al., 2016). However, bLRs, the more anxious and depressive-like phenotype, may be more responsive to drugs that alleviate these symptoms, and are generally more sensitive to the neurobiological impact of aversive stimuli. Since either phenotype could exhibit drug abuse liability, we wanted to assess the impact of adolescent cocaine experience on each phenotype trajectory, measure whether cocaine’s impact during adolescence caused different neurobiological and epigenetic changes in bHRs vs. bLRs, and determine if these changes persist into adulthood and associate with adult psychomotor sensitization, an indicator of responsiveness to psychostimulants.

In particular, bHR rats sensitized both in adolescence and adulthood to this particular repeated cocaine paradigm, and this is consistent with our prior studies (e.g., García-Fuster et al., 2010). However, bHR rats with a prior history of adolescent cocaine, no longer exhibit sensitization in adulthood, suggesting the induction of metaplasticity. The measures presented in this article that best show this phenomenon are locomotor distance, and head-waving frequency. In contrast, although bLRs did not display psychomotor sensitization when given cocaine in either adolescence or adulthood alone (García-Fuster et al., 2010), a prior history of cocaine exposure during adolescence appeared to have “primed” bLRs such that they express increased psychomotor activation later in adulthood following an acute challenge. Thus, the drug experience during adolescence appeared sufficient to override an inherent resistance to expressing psychomotor sensitization in those less addiction liable (i.e., bLRs). However, this acute effect appeared transient, with repeated exposure to cocaine either showing no increase or even a decrease in psychomotor activation on D7 relative to D1.

The fact that identical cocaine treatment regimens in either adolescence or adulthood produced such different behavioral profiles underscore the importance of considering genetic background and associated neurobiological differences in shaping drug responsiveness. The idea that bHRs and bLRs have unique drug responses is consistent with our previous findings showing that an acute cocaine challenge in unmanipulated bHRs and bLRs produces different neurochemical responses in NAc dopamine and norepinephrine levels (Mabrouk et al., 2018). However, the biological mechanisms that shape their contrasting responses to repeated drug exposure across two stages of life are far from understood.

### The Impact of Adolescent and Adult Exposure to Cocaine on the Balance of a Repressive and a Permissive Modified Histone

In this study, we asked whether the differential impact of adolescent cocaine on the bHR vs. bLR adult phenotype is associated with distinct epigenetic changes in the two lines. We used an anatomical, cell-count-based approach to identify epigenetic consequences of adolescent drug experience on subsequent adult drug sensitization in bHR and bLR rats. We focused on specific sub-regions of the striatum and NAc and characterized their profiles by quantifying the levels of the repressive H3K9me3 vs. the permissive acH3K9 at the cellular level. This approach assesses, with single cell granularity, the total change in the level of these histones that can modify the transcription of multiple target genes in those cells. It represents an alternative to techniques such as CHIP-seq that are typically carried out at the level of an entire brain region and aim at identifying the binding sites of a histone mark to specific genes. Focusing on select sub-regions of the striatum we found specific bHR-bLR differences in epigenetic markers, particularly notable in the NAc core.

The two epigenetic marks we examined have been previously shown to be important in cocaine sensitization. Overexpressing permissive acH3K9 produced greater sensitization, while overexpressing similar restrictive di-methylation H3K9 blunted sensitization in mice (Heller et al., 2016). However, these chromatin modifications were directly manipulated on the Cdk5 gene to produce these effects, which is known to modulate DARPP-32 dopamine signaling (Bibb et al., 1999) and phosphorylate D2, regulating its downstream signaling (Jeong et al., 2013). In our model, we have found basal differences in bHR-bLR epigenetic profiles (Chaudhury et al, 2014). In particular, H3K9me3 was found to negatively regulate D2 receptors in the striatum of bHRs following a history of cocaine self-administration (Flagel et al., 2016).

In this study, we found that decreases in H3K9me3 track with the behavioral manifestation of sensitization. In the NAc core of bHRs, H3K9me3 exhibits a pattern consistent with limits on plasticity described at the behavioral level. Thus, its levels are decreased by chronic cocaine in both adolescence and adulthood, and since it is usually repressive, this change is consistent with increased behavioral activation. Similarly in outbred rats, the number of cells expressing repressive H3K9me3 was reduced in the NAc following cocaine sensitization (Maze et al 2011). Moreover, our findings show that the decrease in this histone is stable in that it is seen in saline-treated adults with a history of adolescent cocaine exposure. However, the additional cocaine exposure in adulthood does not further impact to the change in NAc core. Thus, one explanation for why bHRs did not show further augmentation of locomotor sensitization is that the plasticity of their neuroplasticity, or “metaplasticity,” may become diminished by the adolescent cocaine experience, weakening the brain’s response to subsequent drug experience. Alternatively, plasticity might remain intact, but the response takes on a different form to reduce subsequent sensitization to the adult drug exposure (i.e., reduced H3K9me3 in NAc core). Moreover, in the NAc shell of bHRs, there were no cocaine-induced changes in H3K9me3 observed in either adolescents or adults with no prior drug history. Only the double hit exposure (adolescence + adulthood) triggered a change which was in the opposite direction (i.e., towards increased repression), which may contribute to the lack of behavioral sensitization in the Coc-Cocaine group. In bLR rats, there were no changes observed in the expression of this modified histone in the NAc core or shell, in line with their non-sensitized response at the end of the adulthood treatment.

As for the regulation of the permissive mark in bHRs rats, the changes observed in acH3K9 were region-specific (only in NAc core) and showed a consistent pattern with sensitization in both adolescent and adults with no cocaine history. These findings are in agreement with the literature showing that repeated psychostimulant administration increases histone acetylation in the NAc of mice (Kennedy et al., 2013).

The effects on acH3K9 were also persistent, as seen in the Coc-Saline group. However, a double hit cocaine exposure actively reversed this elevation, and decreased the level of this modified histone, which would also be inhibitory (i.e., no further psychomotor sensitization beyond the levels elicited by the first regimen of adolescent cocaine). The change in profile of H3K9me3 and acH3K9 during sensitization and metaplasticity is illustrated in Fig. 5. Overall, if a bHR animal has a dual exposure to cocaine and is showing a limit to further sensitization, several epigenetics changes are consistent with limiting activation or enhancing repression: (a) no further inhibition of H3K9me3 in the NAc core, (b) activation of H3K9me3 in the shell, and (c) inhibition of acH3K9 in the core. All of the changes are consistent with the notion of metaplasticity.

As for bLR rats, acH3K9 was increased in the NAc core, but only after a double hit of cocaine, suggesting it might not be sufficient to cause sensitization in these rats. Interestingly, these changes were restricted to the NAc area, since acH3K9 was not altered in dmSTR or dlSTR of bHR or bLR rats following cocaine treatments. Altogether, the current findings proved that these changes were anatomically (e.g., NAc core vs. shell or STR regions) and histone (methylation vs. acetylation) specific, and were expressed differently in bHRs and bLRs following the same history of adolescent cocaine experience. This suggests these histone modifications are situated to have an active and circuitry-specific regulatory role in the expression of cocaine sensitization, since decreased repressive histone, H3K9me3, and increased permissive histone, acH3K9, associated with cocaine sensitization in the NAc core of bHR rats.

### General Considerations

These findings reflect that adult epigenetic function is both sub-region specific and phenotype specific. The lens of epigenetics offers novel perspectives on how genetics and environment share common mechanisms for ensuring adaptive neurodevelopment, but also the potential for exacerbating maladaptive risk. While we have focused in this study on changes in two modified histones, it is important to note the existence of hundreds of identified posttranslational chromatin modifications which likely interact with several other classes of epigenetic modifications (e.g., DNA methylation, lncRNAs). These modifications likely work in concert as a “histone code” to enact substantive downstream events via transcription factors to drive gene regulation (Strahl and Allis, 2000; Matthews and McGowan, 2019). Thus, vulnerability to psychostimulant use and abuse is likely shaped by broad patterns of gene expression modified by epigenetic and enzymatic modifications in critical neuronal networks that shape responsiveness to the drugs (Lud Cadet, 2016). The advent of technologies such as the use of Transposase Accessible Chromatin with high-throughput sequencing (ATAC-Seq) (Buenrostro et al, 2013) coupled with RNAseq promises to be powerful in the context of drug abuse research for fully characterizing chromatin accessibility, broadly defining epigenetic profiles in a given tissue and cell types to identify which genes are available for tissue-specific transcription. The use of these coupled techniques to understand the interplay between genetic background and adolescent exposure to drugs in modifying vulnerability to drug use or abuse should prove especially powerful.

Our study clearly underscores the importance of adolescence as a critical developmental window for substantial life-long changes via epigenetic modifications that can shape the circuitry for emotion, motivation and addiction. There is growing evidence that epigenetic time windows may accompany critical periods of development, playing a role in the long-term trajectory of the brain (Kanherkar et al., 2014). This may be particularly relevant for addiction liability and outcome given the prevalence of prescribed psychostimulant medication in younger ages and increased accessibility potential for adolescent drug abuse. Epigenetic mechanisms that normally play a functional role in neurodevelopment at that time, including cortical organization, pruning, and coordination of gene expression, may become substantially altered by adolescent drug abuse, causing life-long consequences on subsequent addiction liability.

## Conclusions

We have shown the clear interplay between genetic background and exposure to cocaine during adolescence in altering behavioral phenotypes in adulthood in response to a psychostimulant drug. In particular, we demonstrated that early experience with psychostimulant drugs can limit plasticity in adulthood. Finally, we have demonstrated how epigenetic mechanisms within the NAc, that can affect the broad level of gene activation or repression, are associated with cocaine sensitization and metaplasticity.

## Acknowledgments

We would like to thank Anna Gencay, Jesse Klein, and Natalie Tarrant for their assistance quantifying stereological results, Alex Stefanov for oversight of the selective-breeding colony, and Amy Tan, Fei Li and Jennifer Fitzpatrick for their technical assistance.

## Author Contributions

AP, MJG-F, SBF, and HA designed this research and wrote the paper. AP and MJG-F also performed the research and analyzed data. EH-B and SJW contributed unpublished reagents/analytic tools critical to these experiments.

