## Supplementary materials and figures for "Cocaine during adolescence differentially impacts psychomotor sensitization and epigenetic profiles in adult male rats with divergent affective phenotypes"

### Supplemental Materials and Figures

*Basal Locomotor Response to Novelty – Screening bHR and bLR Phenotypes*

bHR and bLR rats were placed in a novel environment at age P25 (baseline) and total locomotor activity (the sum of horizontal and rearing movements) was measured. Two-way ANOVAs (independent variables: Phenotype and Squad) showed main effects of Phenotype for total locomotion, as well as for horizontal and rearing locomotor activity (see statistical results reported in Supplementary Table S1, and Fig. S1). For all measurements, bHRs showed higher basal activity (i.e., exploratory locomotor behavior in a novel environment) than bLRs. These basal locomotor scores were used to counterbalance bHR and bLR rats to ensure there were no group differences within each phenotype for the subsequent behavioral testing (see individual values per Squads, Supplementary Fig. S1).

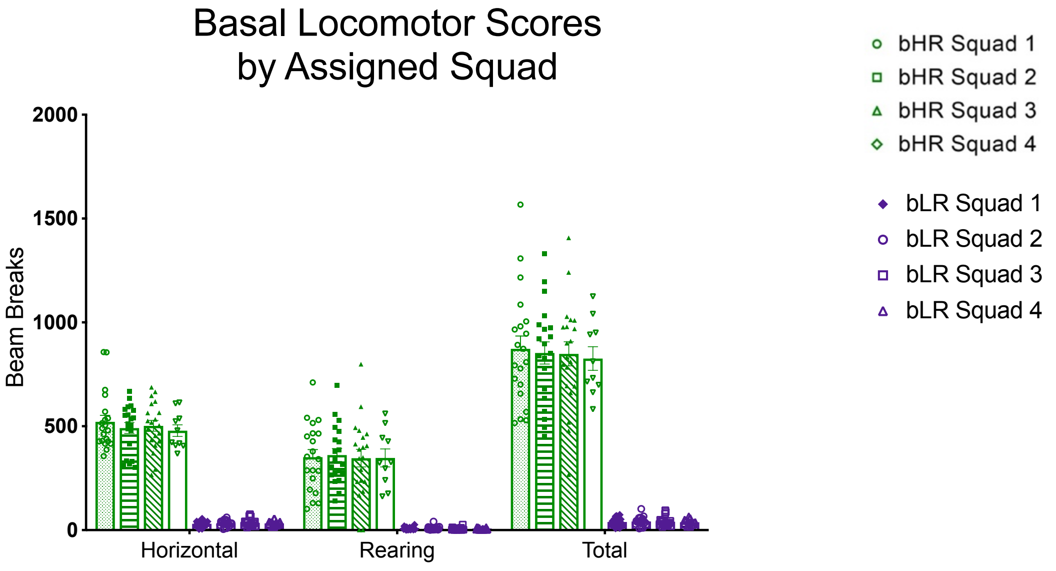


**Fig. S1.** Locomotor response to novelty. bHRs and bLRs predictably differed in their basal locomotor response to novelty. However, all tested conditions were counterbalanced within run squads to ensure that no basal differences biased any subsequent result of testing with cocaine or saline.

*Adolescent Cocaine Differentially Impacts Psychomotor Sensitization in Divergent Affective Phenotypes*

*Effects during adolescence:* Both acute (D1) and chronic (D7) cocaine regimens in adolescent bHR and bLR rats increased locomotor velocity and darting bouts (Fig. S2), as observed by the significant main effects of Treatment reported in Supplementary Table S1.

*Effects during adulthood:* bHR rats exposed to adult cocaine and with a prior history of adolescent cocaine showed similar sensitized locomotor activity levels (i.e., locomotor velocity and darting bouts) on D7 vs. D1 in adulthood, and as compared to rats that only received cocaine in adulthood (Fig. S2), thus indicating that bHR rats that were exposed to cocaine in adolescence do not exhibit further sensitization in adulthood.

Interestingly, in bLR rats, the psychomotor effects of adult cocaine exposure were also observed for locomotor velocity and darting bouts (see main effects of Adult Treatment on Supplementary Table S1 and Fig. S2), but without sings of sensitization (lack of increased responses on D7 vs. D1). However, rats with a prior history of adolescent cocaine, and challenged on D1 in adulthood with acute cocaine showed greater psychomotor activation (e.g., higher darting bouts than bLR rats that received a repeated cocaine treatment during adolescence, Coc-Saline vs. Coc-Cocaine group comparisons in Fig. S2, and Supplementary Table S1).

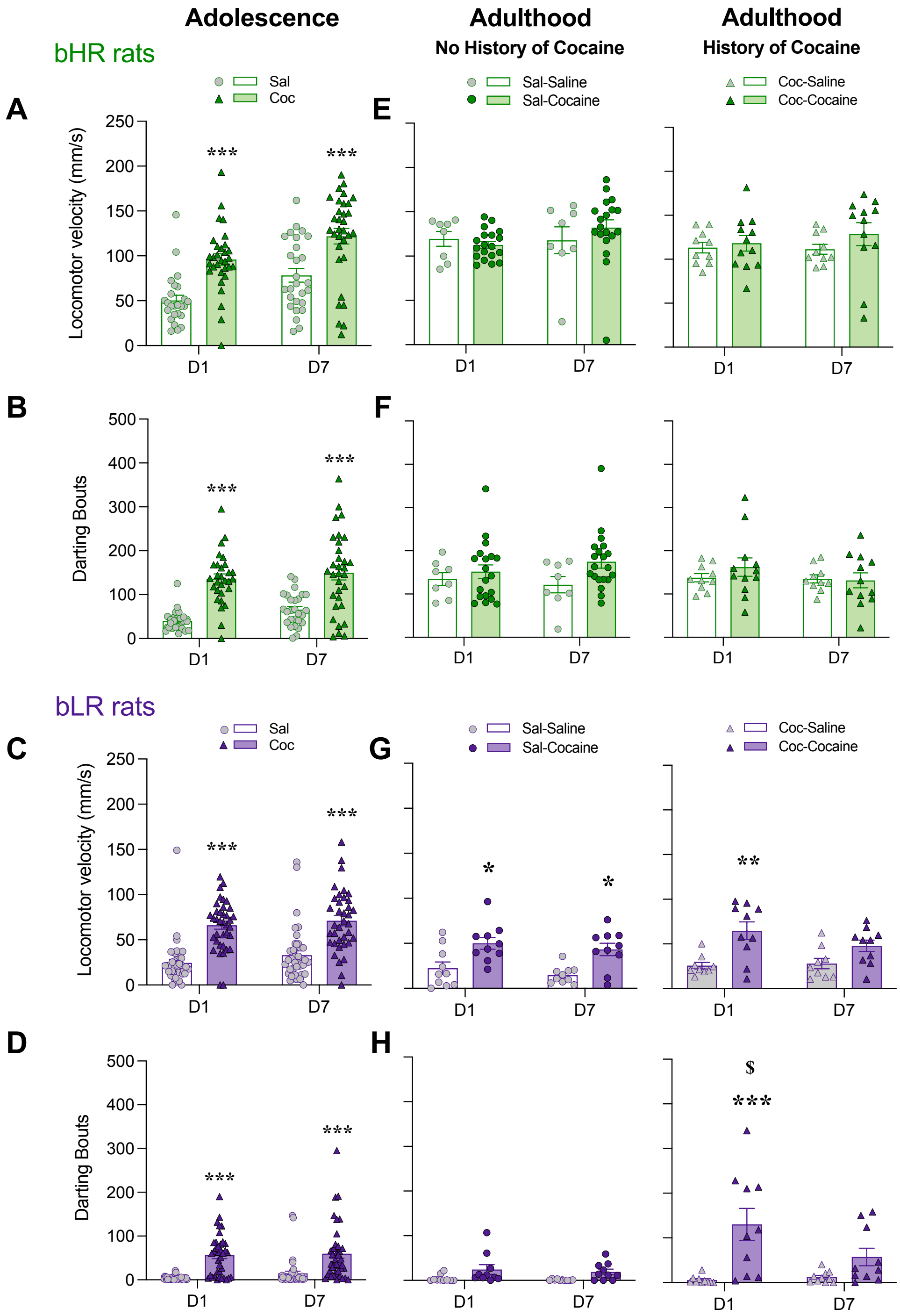


**Fig. S2.** Adolescent Cocaine Differentially Impacts Psychomotor Sensitization in Divergent Affective Phenotypes. Acute (D1) and chronic (D7) psychomotor activating effects of cocaine administration in adolescent (panels A-D) and/or in adulthood (with no or with a prior history of adolescent cocaine; panels E-H) in bHR and bLR rats. Data represents the mean values ± SEM of the locomotor velocity (mm/s) (panels A, C, E and G) and the number of darting bouts (panels B, D, F and H). Individual values are shown for each rat (symbols). Two- or three-way RM ANOVAs were used for statistical analysis (see Supplementary Table S1 for more details). $*p < 0.05$, $**p < 0.01$ and $***p < 0.001$ when comparing cocaine-treated rats with the corresponding saline-treated group (main Effect of Treatment). $\$p < 0.001$ when comparing the response on D7 of the adolescent treatment vs. the one following an acute challenge dose in adulthood (D1 for Coc-Cocaine group).

*Baseline levels of H3K9me3 and acH3K9 expression in the NAc of bHR and bLR rats*

There were no baseline differences between bHRs and bLRs control rats (saline-treated) in expression of either H3K9ME3 or acH3K9 histone mark (Fig. S2) with age (adolescence, P40 vs. adulthood, P85).

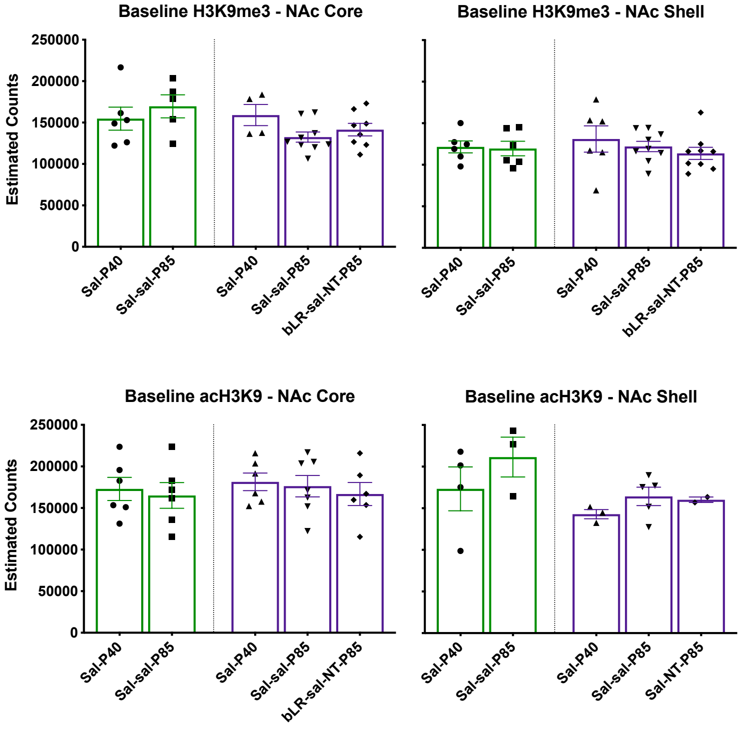


**Fig. S3.** bHR (green) and bLR (red) baseline NAc differences in H3K9me3 and acH3K9. No basal differences were seen in either marker among control-treated rats and when comparing adolescence with adulthood expression.

*Adolescent Cocaine Did Not Impact the Expression of acH3K9 in the Dorsomedial and Dorsolateral Striatum of Divergent Affective Phenotypes*

We also quantified the number of cells expressing acH3K9 in the dorsomedial (dmSTR) and dorsolateral (dlSTR) regions of the striatum in adult bHR and bLR rats (Fig. S4). There were no significant changes induced by cocaine exposure (adolescent, adult or both) in these subregions in either bHR or bLR rats (Fig. S4; Table S1).

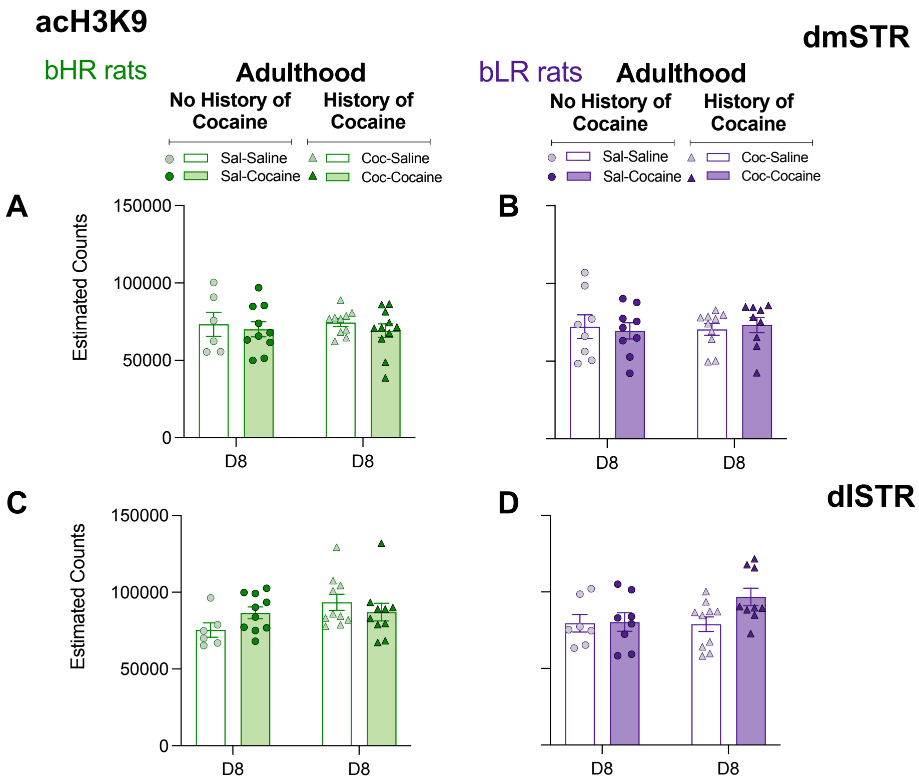


**Fig. S4. Adolescent Cocaine Did Not Impact Epigenetic Expression in the Striatum (dorsomedial, dmSTR and dorsolateral, dlSTR) of Divergent Affective Phenotypes.** Permissive acH3K9 (panels A-D) cell counts in the dmSTR and dlSTR of bHRs and bLRs treated with adolescent and/or adult cocaine. Data represents the mean values ± SEM of the estimated cell counts of acH3K9 as measured 24 h post-adult treatment (D8). Individual values are shown for each rat (symbols). Two-way ANOVAs were utilized following the adult treatment (see Supplementary Table S1 for more details) and showed no significant differences.
