## Supplementary figures and images for "Cocaine during adolescence differentially impacts psychomotor sensitization and epigenetic profiles in adult male rats with divergent affective phenotypes"

### Supplementary Table S1

#### Slide 1
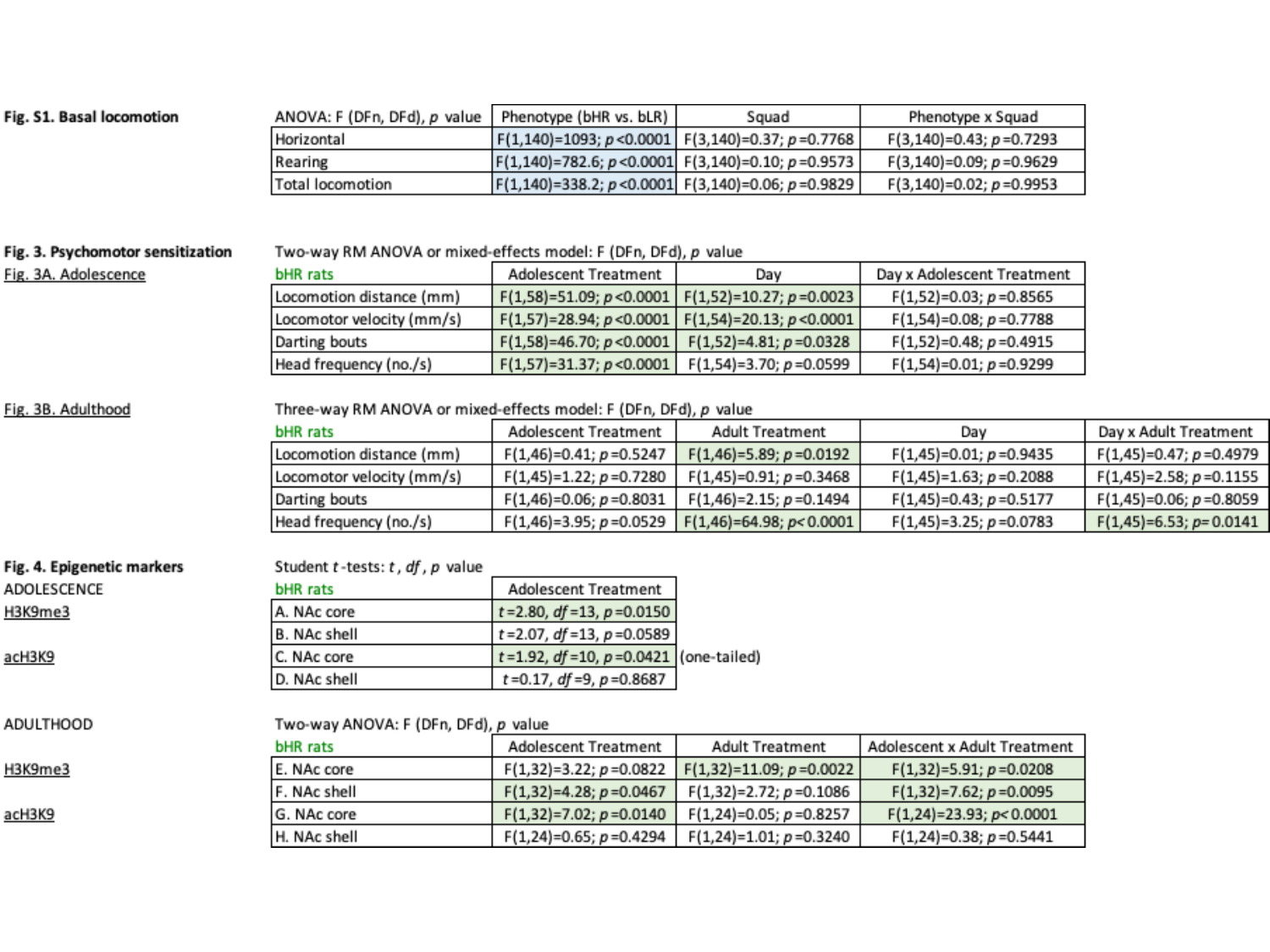

#### Slide 2
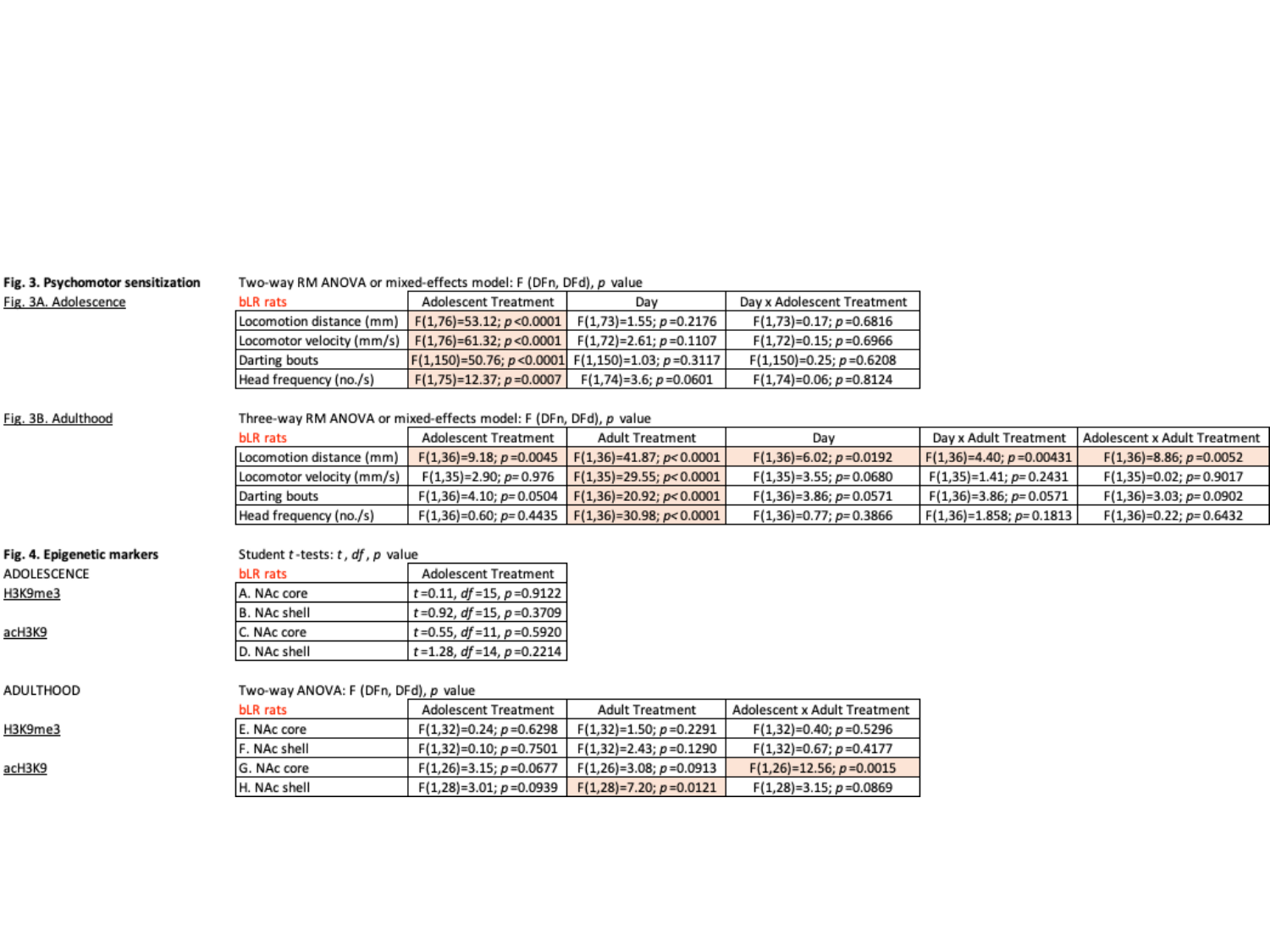
